# Using deep neural networks to detect complex spikes of cerebellar Purkinje Cells

**DOI:** 10.1101/600536

**Authors:** Akshay Markanday, Joachim Bellet, Marie E. Bellet, Ziad M. Hafed, Peter Thier

## Abstract

One of the most powerful excitatory synapses in the entire brain is formed by cerebellar climbing fibers, originating from neurons in the inferior olive, that wrap around the proximal dendrites of cerebellar Purkinje cells. The activation of a single olivary neuron is capable of generating a large electrical event, called “complex spike”, at the level of the postsynaptic Purkinje cell, comprising of a fast initial spike of large amplitude followed by a slow polyphasic tail of small amplitude spikelets. Several ideas discussing the role of the cerebellum in motor control are centered on these complex spike events. However, these events are extremely rare, only occurring 1-2 times per second. As a result, drawing conclusions about their functional role has been very challenging, as even few errors in their detection may change the result. Since standard spike sorting approaches cannot fully handle the polyphasic shape of complex spike waveforms, the only safe way to avoid omissions and false detections has been to rely on visual inspection of long traces of Purkinje cell recordings by experts. Here we present a supervised deep learning algorithm for rapidly and reliably detecting complex spikes as an alternative to tedious visual inspection. Our algorithm, utilizing both action potential and local field potential signals, not only detects complex spike events much faster than human experts, but it also excavates key features of complex spike morphology with a performance comparable to that of such experts.

**Significance statement:** Climbing fiber driven “complex spikes”, fired at perplexingly low rates, are known to play a crucial role in cerebellum-based motor control. Careful interpretations of these spikes require researchers to manually detect them, since conventional online or offline spike sorting algorithms (optimized for analyzing the much more frequent “simple spikes”) cannot be fully trusted. Here, we present a deep learning approach for identifying complex spikes, which is trained on local field and action potential recordings from cerebellar Purkinje cells. Our algorithm successfully identifies complex spikes, along with additional relevant neurophysiological features, with an accuracy level matching that of human experts, yet with very little time expenditure.

## Introduction

The Purkinje cell (PC) output, the sole output of the cerebellar cortex, is characterized by two distinct types of responses (Fig. 1A, bottom), the simple spike (SS) and the complex spike (CS) (Thach, 1968). SSs are ordinary sodium-potassium spikes with a simple bi-or tri-phasic shape in extracellular recordings (Fig. 1B). These spikes, lasting only a fraction of a millisecond and firing up to several hundred times per second, reflect the concerted impact of mossy fiber input, mediated via the granule cell-parallel fiber system, as well as inhibitory interneurons. On the other hand, an individual CS (Fig. 1C), elicited by a single climbing fiber originating from the inferior olivary nucleus and pervading the proximal dendrites of a PC, is characterized by a polyphasic somatic spike consisting of a first back propagated axonal spike component followed by a series of spikelets riding on a long-lasting, calcium dependent depolarization (Eccles et al., 1967; Fujita, 1968; Thach, 1968; Llinas and Sugimori, 1980; Stuart and Häusser, 1994; Davie et al., 2008). In addition to an exceptional morphology, CSs also exhibit an unusual, perplexingly low firing rate of at most two spikes per second (Fig. 1A, bottom). What could these infrequent, yet unique events possibly tell us about their purpose, and what might be the best statistical tool allowing us to unravel the full extent of information carried by them? These are questions that have kept researchers busy until today.

**Figure 1.**
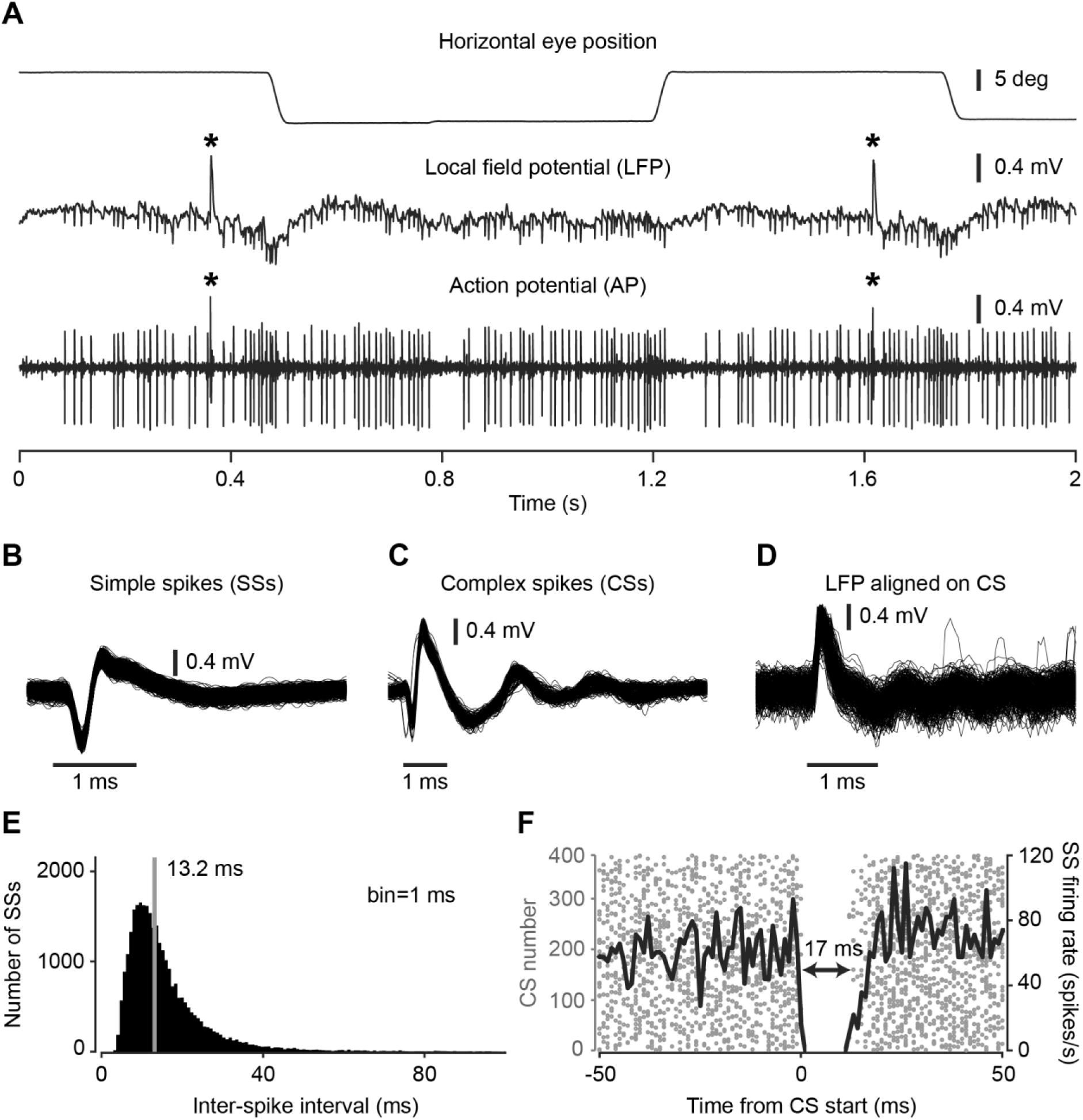
Characteristics of an exemplary Purkinje cell. (A) Local field potential (LFP, low passed, <150 Hz, middle panel) and action potential (AP, high band-passed, 300 Hz - 3 KHz, bottom panel) activity in relation to horizontal eye movements (top panel). CSs are marked by asterisks. (B) Isolated SS waveforms aligned on SS start. (C) Isolated CS waveforms aligned on CS start. (D) LFP responses aligned to CS start. (E) Histogram of inter-spike intervals of SSs. Solid gray line depicts the median value (13.2 ms). (F) Raster plot showing a 17 ms pause in SS activity caused by the occurrence of a CS. Solid black line represents the mean SS firing rate aligned to CS start.

Thinking about the role of CSs has been guided by two, not necessarily incompatible, ideas: motor timing and motor learning. The first idea, championed by Llinás and his coworkers, was prompted by the characteristic 8-10 Hz rhythmicity and synchronicity of inferior olivary neurons, a pattern that seemed to reflect the temporal structure of many forms of motor behavior, as well as physiological and pathological tremor (Llinas, 1974; Leznik and Llinás, 2005). The second idea emphasized the role of performance errors in driving motor learning. On experiencing an error, the climbing fiber system is assumed to produce a CS, which helps to predictively correct future manifestations of the same motor behavior by modifying the impact of parallel fibers on targeting PCs (Marr, 1969; Albus, 1971; Ito, 1972). This concept has indeed received support from a number of experimental studies (Oscarsson, 1980; Kitazawa et al., 1998; Medina and Lisberger, 2008; Herzfeld et al., 2015, 2018). However, not all findings have been fully compatible with this so-called Marr-Albus-Ito hypothesis, at least not in its original form. For instance, recent work on oculomotor learning has suggested that CS discharge is not only influenced by a current error, but also by a memory of past errors suitable to stabilize behavioral adaptations (Catz et al., 2005; Dash et al., 2010; Junker et al., 2018). An analogous influence of past errors on CS discharge has also been noted in recent studies of eye-blink conditioning (Ohmae and Medina, 2015). Finally, others have advocated that CSs may not be confined to encoding unexpected errors, but to also offer a prediction of the multiple kinematic parameters of the upcoming movement (Streng et al., 2017).

Reaching consensus on the diverse views of CS functions would be substantially facilitated by more data on these sparse neural events, collected in conjunction with advanced behavioral paradigms. Yet, it is exactly their unique properties of rarity and complex and highly idiosyncratic spike morphology that have hampered progress. In fact, CS spike morphology not only differs between individual PCs, but it also often changes over the course of a single recording from the same PC. This is why using standard spike sorting software to detect CSs has turned out to be error prone. Critically, given the rarity of CSs, even a few missing or erroneously detected CS events will have profound impacts on conclusions drawn about their functional role. Consequently, researchers are compelled to meticulously label CSs manually, or at least to visually control the CSs detected by conventional spike sorting approaches, an exhausting approach that constrains the amount of experimental data that can be processed.

In this paper, we exploited a state-of-the-art convolutional neural network (CNN) approach to dramatically reduce the burden of investigators in identifying CSs. We show that our network is able to learn fast and that it easily matches the performance of an experienced human expert in detecting CSs. Our algorithm also extracts a number of key parameters on CS timing and morphology, in a regularized and systematic manner, which we believe is particularly important for understanding the functional role of CSs.

## Materials and Methods

### Animals, preparation, surgical procedures, and recording methods

Two adult male rhesus macaques (*Macaca mulatta*) of age 10 (monkey K) and 8 (monkey E) years, purchased from the German Primate Center, Göttingen, were subjects in this study. Initial training of all animals required them to voluntarily enter an individually customized primate chair and get accustomed to the setup environment, a procedure that could last for up to three months. Following initial training, they underwent the first major surgical procedure in which foundations of all implants were fixed to the skull using titanium bone screws, and then allowed to rest for a period of approximately 3-4 months to improve the long-term stability of the implant foundations. Then, a titanium-based hexagonal tube-shaped head post was attached to the implanted head holder base to painlessly immobilize the head during experiments, and scleral search coils were implanted to record eye positions using electromagnetic induction (Judge et al., 1980; Bechert and Koenig, 1996). Within 2-3 weeks of recovery from the eye-coil implantation procedure, monkeys quickly recapitulated the already learned chair-training protocol, and were trained further on their respective behavioral paradigms. Once fully trained, a cylindrical titanium recording chamber, whose position and orientation were carefully planned based on pre-surgical MRI and later confirmed by post-surgical MRI, was finally mounted on the implanted chamber base, tilting backwards by an angle of 30° with respect to the frontal plane, right above the midline of the cerebellum. A part of the skull within the chamber was removed to allow precise electrode access to our region of interest, the oculomotor vermis (OMV, lobuli VIc/VIIa), for electrophysiological recordings. All surgical procedures were carried out under aseptic conditions using general anesthesia, and post-surgical analgesics were delivered until full recovery. See Prsa et al. (2009) for full details. All experiments and surgical procedures were approved by the local animal care authority (Regierungspräsidium Tübingen) and complied with German and European law as well as the National Institutes of Health’s *Guide for the Care and Use of Laboratory Animals.* All procedures were carefully monitored by the veterinary service of Tübingen University.

### Behavioral tasks

In-house software (NREC), running on a Linux PC (http://nrec.neurologie.uni-tuebingen.de), was used for data collection, stimulus presentation, and operations control. The two monkeys were trained on a fatigue inducing repetitive fast eye movements (saccades) task (Fig. 1A, top; Prsa et al., 2010). A trial started with a red fixation dot (diameter: 0.2°) displayed at the center of a CRT monitor placed 38 cm in front of the monkey. After a short and variable fixation period (400-600 ms from trial onset), the fixation dot disappeared and at the same time, a target, having the same features as the fixation dot, appeared on the horizontal axis at an eccentricity of 15°. In a given session, the target was presented consistently either on the left or right of the central fixation dot. The maximum number of trials (>200) per session depended on the willingness of the monkey to cooperate and on the duration for which a PC could be kept well isolated. Each trial lasted for 1200 ms, and inter-trial intervals were kept very short (100 ms) to maximize the induction of fatigue. At the end of every correct trial, monkeys were rewarded with a drop of water.

### Electrophysiological recordings

Extracellular recordings with commercially available glass-coated tungsten microelectrodes (impedance: 1-2 MΩ; Alpha Omega Engineering, Nazareth, Israel) were performed using a modular multi-electrode manipulator (Electrode Positioning System and Multi-Channel Processor, Alpha Omega Engineering) whose position was estimated, based on the position and orientation of the chamber relative to the brain, using a stereotactic apparatus and later confirmed by post-surgical MRI scans. Saccade-related modulation of an intense background activity, reflecting multi-unit granule cell activity, paralleled by saccade-related modulation in the local field potential record (LFP, <150 Hz bandwidth) served as electrophysiological criteria for identifying the OMV (Fig. 1A, middle). Extracellular potentials, sampled at 25 KHz, were high band-pass (300 Hz - 3 KHz) and low-pass filtered (<150 Hz) to differentiate PC action potentials and LFP signals, respectively (Fig. 1A, bottom).

### Multi Spike Detector: the online spike sorting algorithm

Single PC units were identified online by the presence of a high-frequency SS discharge accompanied by the signatory, low-frequency CS discharge using a real-time spike sorter, the Alpha Omega Engineering Multi Spike Detector (MSD). The MSD, designed for detecting sharp waveforms uses a template matching algorithm developed by Wörgötter et al. (1986), sorts waveforms according to their shape. The algorithm employs a continuous comparison of the electrode signal against an 8-point template defined by the experimenter to approximate the shape of the spike of interest. The sum of squares of the difference between template and electrode signal is used as a statistical criterion for the goodness of fit. Whenever the goodness of fit crosses a threshold, the detection of a spike is reported. The 8-point template can be adjusted manually or alternatively, run in an adaptive mode that allows it to keep track of waveforms that may gradually change over time.

### Identification of simple spikes and complex spikes in Purkinje cells

As opposed to short duration SSs (Fig. 1B), characterized by short median inter-spike intervals (Fig. 1E), the long duration CSs (Fig. 1C) were much more rare. In addition to the 10-20 msec long pause triggered by a CS in the SS firing (e.g. Fig. 1F, Bell and Grimm, 1969; Latham and Paul, 1971; McDevitt et al., 1982), the presence of a CS was also indicated by a massive deflection of the LFP signal, lasting for the whole duration of a CS (Fig. 1D). While the MSD-based detection of abundantly available SS events can be trusted most of the time, since the consequences of erroneously including or missing a few SSs are less problematic, MSD-based detection of much rarer CS events is error prone, the costs of which cannot be neglected. Consequently, thorough analysis of PC data often requires experimenters to visually control the quality of MSD-based detections post-hoc, and many times, to even manually identify CS events.

### Convolutional neural network

We used the architecture of a CNN that was originally designed to segment images (“U-Net”, Ronneberger et al., 2015) and later successfully adapted for the detection of saccades in eye position recordings (“U’n’Eye”; see Bellet et al. (2018) for details). For CS detection, we input the LFP and action potential signals, sampled at the same frequency of 25 KHz, to the network (Fig. 2A, top). The output was a bin-wise predictive probability of CS occurrence (Fig. 2A, bottom).

**Figure 2.**
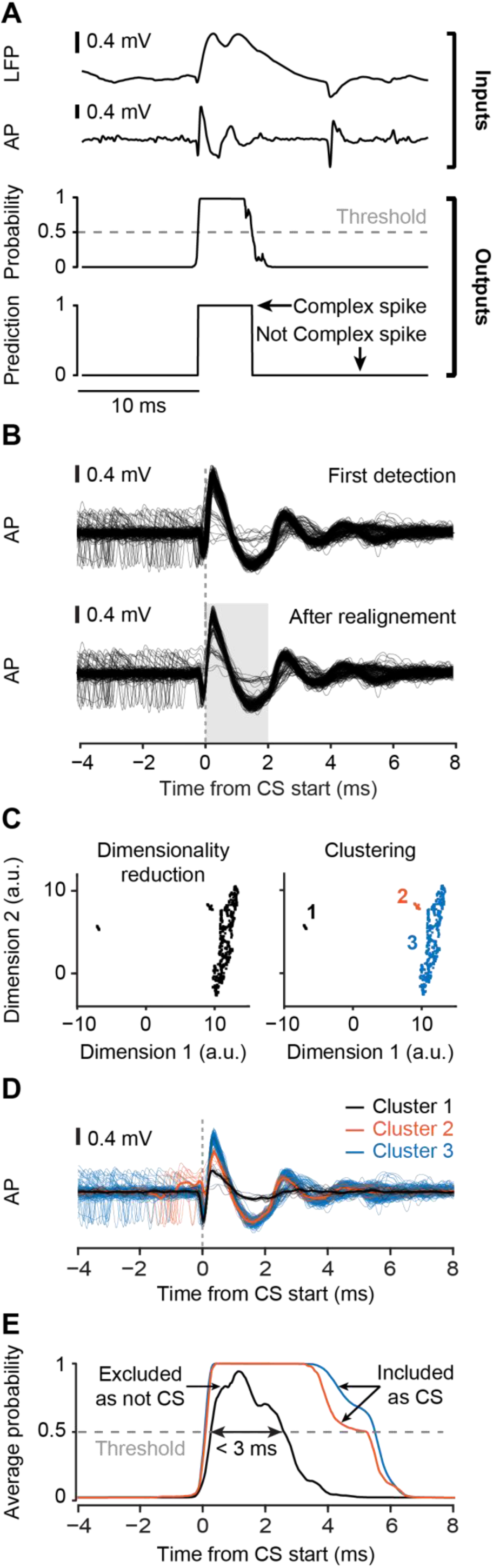
Pipeline for complex spike detection. (A) Input to the network (LFP and action potential signal, labels as AP) as well as its output (bin-wise predictive probability for CS occurrence and binary CS classification). (B) Waveforms aligned to the first estimation of start times of all CSs detected by the network (upper panel) used for computing an average waveform that served as a template for realigning the waveforms of all detected CS events (lower panel). (C) Projection of the waveforms during the time interval shaded in gray in B onto a two-dimensional plane and identification of clusters in this space. Different colors indicate distinct clusters. (D) Waveforms of the clusters in (C). Note that Cluster 1 clearly violates well-known CS waveform shapes. (E) Average predictive probability output of the network for the events in each cluster. Clusters, whose probability output exceeds the classification threshold of 0.5 (dashed gray line) for less than 3 ms, are excluded as not representing CSs (Cluster 1).

The network consists of convolutional and max-pooling layers. Max-pooling is an operation that down-samples the input in order to reduce the dimensionality of its representation in the network. It filters the input with a certain window size and extracts only the maximum value. It then steps further on the input, repeating the same operation on the next time window. Convolutional layers extract relevant features of the input signal by learning the parameters of its convolutional kernel during training. We chose the size of the max-pooling (mp) and convolutional kernels (c) as 7 and 9 bins, respectively. These influence the signal interval (SI) taken into account for labeling one time bin in the output, as described by the formula,

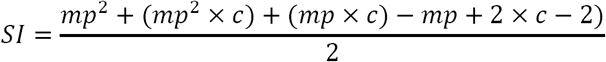

In our case, the SI corresponds to 281 time bins before and after each classified bin.

### Training and testing procedures

We recorded a total of 160 PCs, out of which 119 PCs were selected, based on careful visual assessment of MSD-based CS detection by a human expert (author AM), for in-depth statistical analysis. These PCs remained stable throughout the recording session with clearly isolated CSs and associated signatory SS pauses and LFP deflections. The remaining 41 PCs, for which it was deemed that MSD-based analysis might have led to spurious detections of SSs and CSs, were excluded from analysis.

To prepare the training set, we asked our human expert, who is experienced in electrophysiological recordings from PCs, to visually identify CS events and manually label their start and end points. The expert used small segments of action potential and LFP recordings during labeling, without access to eye movement data. For each PC, 24 segments, each 250 ms long, were manually labeled. To avoid having segments in which a part of a CS may have been truncated (at the beginning or end of a segment), we excluded the first and last 9 ms of each segment during training, thereby reducing its size to 232 ms. Since the network was trained on the manually labeled data, recording segments from the excluded set of 41 PCs, for which the MSD-based CS detection was poor but the human expert-based visual identification was still feasible, were also included for training the network in addition to the selected set of 119 PCs. The number of recording segments for a given PC included in training naturally varied with the number of CSs found in the particular cell, but we ensured including recording segments from all 160 PCs in training.

Since the MSD-based CS detection in 41 PCs was already unsatisfactory, as stated above, a comparison based on the performance of our algorithm and the MSD on these particular PCs would have been too biased in favor of our algorithm. Therefore, to fully test our algorithm’s performance while still giving the MSD-based approach the benefit of the doubt, we used cross-validation on recordings from only the selected pool of 119 PCs. For every PC tested for CS detection, we trained a separate network excluding the currently tested PC from the training set. This allowed us to test how well the network generalized to new data sets, on which it had not been trained, and it also allowed us to have multiple performance tests on our algorithm. Therefore, the training set always comprised the remaining 159 PCs not being currently tested. The total number of recording segments used in any given training set was 970-988, depending on the PC under test. Other parameters of network training such as loss function, learning rate, batch size, and early stopping criterion, were chosen as described in Bellet et al. 2018 for U’n’Eye.

We also performed one more performance test of our algorithm, which was concerned with establishing consistency with expert labeling. For 7 PCs (out of our 119 selected ones described above), we asked our human expert to manually label CSs in the entire records, and not just a small training subset within each of them. This allowed us to directly compare the labeling of the entire records of these 7 PCs by both our algorithm and the human expert. Our algorithm in this case was based on training the network on segments from the remaining 159 PCs (other than the currently tested one), as described above.

### Post-processing

We implemented three post-processing steps to enhance the quality of CSs detected by our algorithm. First, time shifts between the detected start points of all CSs fired by a particular PC were corrected by re-aligning them. To this end, we computed the average waveform from the first estimation of start times of all detected CSs. This average-waveform template was then used as a reference to realign each waveform within a ±2 ms window around CS start so that the cross-correlation was maximized (Fig. 2B). Second, action potential and LFP waveforms, occurring within 2 ms after CS start, were projected onto a two-dimensional plane (Fig. 2C) using the UMAP dimensionality reduction technique (McInnes et al., 2018). This allowed us to use the third post-processing step to cluster waveforms into suitable CSs and unsuitable ones. In this third step, groups of waveforms were identified (Fig. 2D) using HDBSCAN, a hierarchical clustering algorithm (Campello et al., 2013) that builds a tree to describe the distance between data points. The algorithm minimizes the spanning size of the tree and further reduces the complexity of the tree to end up with a minimum number of leaf nodes, corresponding to the clusters. We used the default parameters for HDBSCAN with the option to find only one cluster. Waveforms were excluded if they belonged to a cluster for which the average predictive probability output from the network remained below 0.5 for more than 3 ms (Fig. 2E).

### Quality metrics

We evaluated the performance of our algorithm in detecting CSs using the so-called F1 score (Dice, 1945; Sørensen, 1948), which compares the consistency of CS labels predicted by the algorithm, to “ground-truth” labels provided by the human expert. The F1 score is the harmonic mean of recall (the ratio of true positive detections and all true CS labels) and precision (the ratio of true positive detections and all CS labels predicted by the algorithm), as given by the following equation

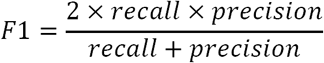

In our case, an F1 score of 1 would suggest that the CSs predicted by our algorithm perfectly matched the “ground-truth” labels provided by the human expert. However, a lower F1 score may suggest that CSs were either erroneously missed or falsely detected. For quality assessment, we also computed the post-CS firing rate of SSs, a signatory feature immune to labels detected by the human expert, which served as a reliable and objective criterion for the identification of a CS. Finally, the resulting CS waveforms were scrutinized by visual inspection.

## Results

### CNN-based algorithm reliably detects complex spikes

The main idea of our approach was to train a classifier to extract relevant features from electrophysiological recordings of PCs and to identify CSs. This was realized with the help of a CNN that uses the LFP and action potential signals as inputs (Fig. 2A, top). We chose these two inputs because human experts achieve consensus on the presence or absence of a CS, more easily and reliably, if both action potentials and LFPs are simultaneously available. Our network uses convolutional and max pooling operations to extract the temporal features relevant for distinguishing CSs from the surrounding signal. In the end, the network predicts the probability of the presence of a CS for each time bin. Time bins for which the predictive probability exceeded the threshold of 0.5 are classified as CSs (Fig. 2A, bottom). The prediction for each time bin depends on an interval in the input signal whose size is determined by the size of the max-pooling and convolutional kernels of the CNN (Methods). Our analysis considered an interval of 281 time bins before and after the time bin containing a predicted CS event. As our sampling rate was 25 kHz, a 10 ms duration CS would span 250 time bins. This means that the network was often using information surrounding CS events (281 versus 250 time bins) to classify CSs.

One of the key requirements for correct CS classification is the quality of the recorded PC signal, which may naturally depend on several factors. For example, subtle drifts between electrode tip and the cell body during a recording session can lead to sudden or gradual changes in the signal-to-noise ratio of the PC signal, and potentially change the morphology of the CS waveform. Also, several SSs firing in close proximity to each other might lead to complex waveforms that may erroneously be detected as CS events. Furthermore, there is also a possibility of CS waveforms being modified by the presence of preceding SSs (Servais et al., 2004; Zang et al., 2018). In order to make our algorithm more resilient to such influences, we added automatic post-processing steps at the output of the CNN. We first fine-tuned the CS start points (Fig. 2B, Methods), and we then differentiated between candidate waveforms using a clustering algorithm in a dimensionally-reduced space (Fig. 2C, Methods). The waveform clusters after dimensionality reduction represented potential candidates for CSs of the recorded PC. Some of these candidates needed to be excluded. For example, if the network in the first step mistakenly classified non-CS events as CSs, then the clustering method would help to refine the classification and exclude these events post-hoc: amongst the CS events erroneously detected by the network might be SSs that are revealed by a separate cluster in the two-dimensional space (Fig, 2C and D, black vs. orange and blue). These false positive events were removed by applying a threshold to the average predictive probability output of the network of the respective cluster (Fig. 2E). Not only non-CS events might have contributed to a distinct cluster separated from the main CS cluster, but true CSs with slightly deviant waveforms (Fig. 2D orange vs. blue) might also have led to separate clusters in the two-dimensional space (Fig 2C orange vs. blue). For all CS clusters that met the defined threshold criterion on predictive probability (Fig. 2E, cluster 1 and 2), the output of our algorithm, CS timing and corresponding cluster IDs, allowed the user to carefully inspect each cluster and decide whether to include clusters with deviant, yet true, CSs or not.

### Objective quality measure confirms identity of complex spikes

It is well-established that SS firing rate decreases during 10-20 ms after the emission of a CS (Bell and Grimm, 1969; Latham and Paul, 1971; McDevitt et al., 1982, Fig. 1F). This physiological feature, independent of the subjective assessment of the human expert, provided us with an additional means for objectively measuring the CS labeling quality of our algorithm. For 119 PCs, we evaluated SS firing rates before and after the occurrence of CSs detected by our algorithm. As depicted in Fig. 3, CSs identified by the algorithm were followed by a clear and significant decrease in the neurons’ SS firing rates by 96% on average (Fig. 3A). In the pre-CS period of 3 to 8 ms, median SS firing rate of the 119 PCs was 58.7 spikes/s; this dropped to 10.5 spikes/s in the post-CS period of 10-15 ms (Fig. 3B, Wilcoxon signed-rank test: p = 2.18 × 10^−20^). This indicates a very low probability of false positive CS detections, since such false positives would increase the apparent post-CS firing rate of SSs.

**Figure 3.**
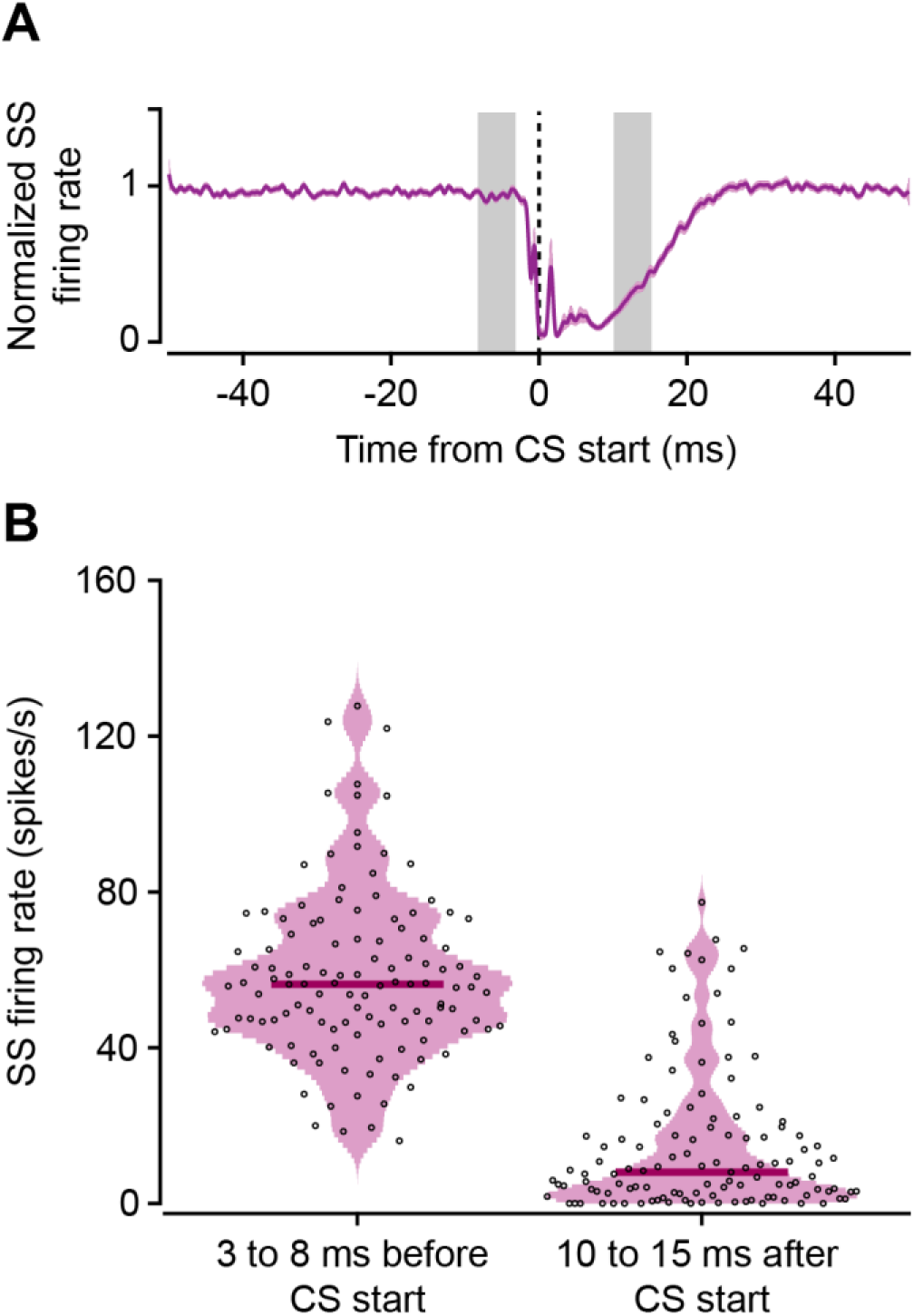
Decrease of SS rate after CSs. (A) Baseline-normalized mean SS firing rate aligned to the start of CSs detected by our algorithm. Data shows mean ± SEM over 119 PCs. Note that the small sharp peak in the SS response, seen immediately after CS start (vertical dashed line in black), is a result of the detection of initial large components of CSs in some PCs where these initial components resembled the shape of SSs and were most probably falsely detected as SSs by the online sorter. (B) Violin plots showing SS firing rate -8 to -3 ms before and 10 to 15 ms after CS start. Each dot represents the average SS firing rate aligned to start time of all CSs in one PC predicted by our algorithm. Thick lines indicate the median SS firing rate of all PCs.

### The new algorithm outperforms a widely-used online sorter

The spike sorting application MSD, based on a template matching algorithm suggested by Wörgötter et al. (1986) for online CS detection, has been widely used by several laboratories as an aid in supporting the visual inspection of PC records (e.g. Catz et al., 2005). This is why we compared the performance of our CNN-based approach to that of the MSD for the same 119 PCs used to test the performance of the algorithm in the previous section. Overall, our algorithm detected 23% more CS events than the MSD (p = 1.4 × 10^−25^, Wilcoxon signed-rank test; Fig. 4A). In order to objectively quantify the difference in CS detection by our algorithm and the MSD, and to verify that the additionally detected events were indeed CSs, we again evaluated the decrease of post-CS SS firing rate. The median decrease of SS firing rate after CSs detected only by our algorithm and not by the MSD was significantly stronger than the decrease induced by CSs detected only by the MSD and not by our algorithm (p = 1.4× 10^−5^, Wilcoxon signed-rank test; Fig. 4B). This indicates that the CSs detected by our algorithm and missed by the MSD were veridical, whereas CSs only detected by the MSD and not by our algorithm were probably erroneous detections (false positives). This view is also supported by a consideration of the time course of SS firing rate aligned to the start time of detected CSs. SS firing rate for CSs only detected by our algorithm and not by the MSD revealed a peak, approximately 3 ms earlier than in the case of CSs that were detected only by the MSD (Fig. 4C). This suggests that SSs occurring shortly before a CS altered the waveform of the latter (Servais et al., 2004) (also see Fig. 2D showing how the amplitude of the average CS waveform of cluster 2 was reduced), therefore impeding its detection by the MSD.

**Figure 4.**
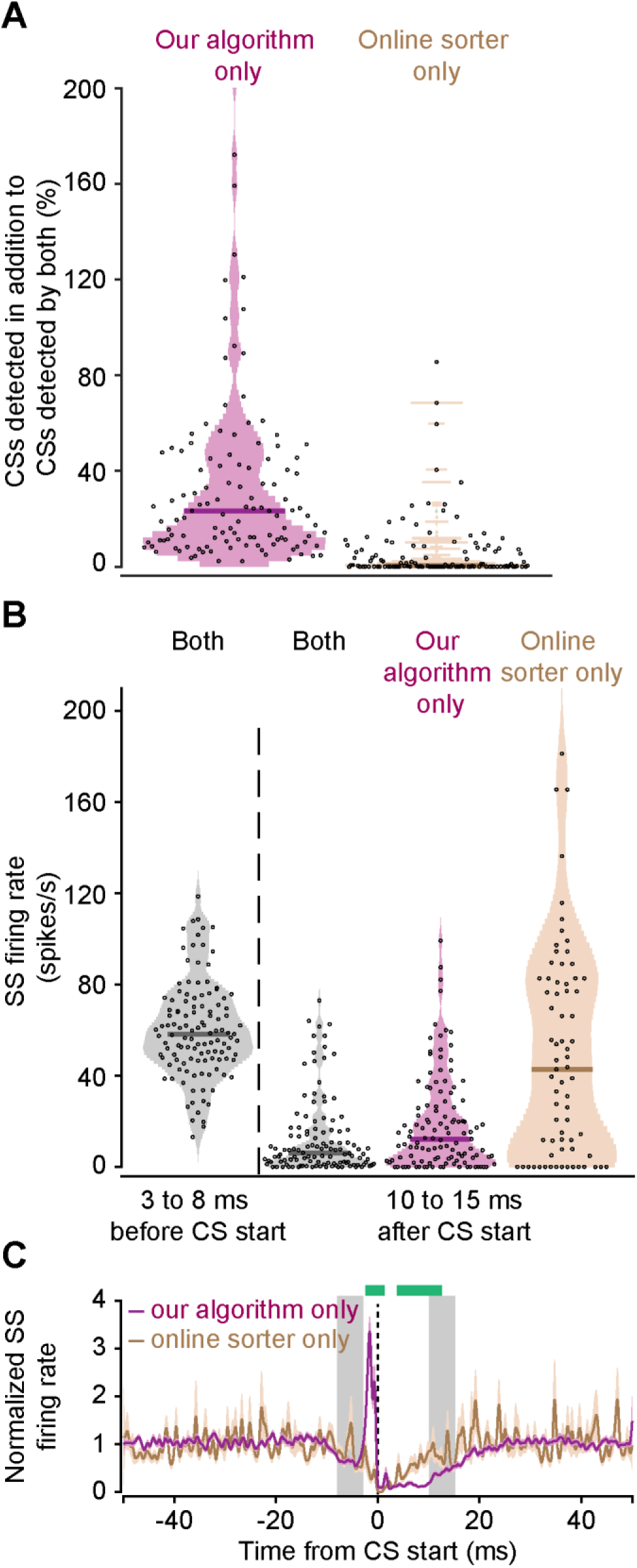
Comparison of CS detection by our algorithm and by the online sorter application, MSD. (A) Violin plots showing percentage of CSs detected exclusively by our algorithm and the online sorter. 100% corresponds to the number of CSs detected by both methods. Our algorithm detected significantly more CSs than the MSD. (B) Violin plots showing SS firing rate aligned to the start of the CSs predicted by both algorithms (gray) or of the events additionally labeled as CSs by either our algorithm (pink) or the online sorter (beige). The decrease in SS firing after CSs predicted by our algorithm but not by the online sorter indicates a higher sensitivity of our algorithm. (A and B) Each dot represents the average SS firing rate aligned to all CSs for the recording of one neuron. Thick lines indicate the median. (C) Pause in averaged SS firing rate following a CS. Gray shaded region represents the period of 3-8 ms before and 10-15 ms after CS start used for comparing SS firing rates in panel B. The sharp increase in SS firing rate approximately 3 ms prior to CS start (vertical dashed line in black), observed only for CSs detected by our algorithm (pink), and not the MSD (beige), suggests that these SSs occurring shortly before the start of CSs might have altered their waveform. Only our algorithm was sensitive enough to detect such CSs with altered waveforms. Green bars on top show intervals with a significant difference between the two traces (random permutations cluster-corrected for multiple comparisons).

We also found that CS waveforms for CSs only detected by our algorithm and not by the MSD were similar in shape to the CSs detected by both our algorithm and the MSD (Fig. 5, middle column vs. left). CSs labeled only by the MSD, on the other hand, deviated from this waveform shape (Fig. 5 right vs. left). This impression clearly also concurs with the weaker post-CS depression of SS firing rate seen in the pool of CS events detected only by the MSD (Fig. 4C). In summary, our algorithm is both more sensitive and less error prone than the MSD-based detection.

**Figure 5.**
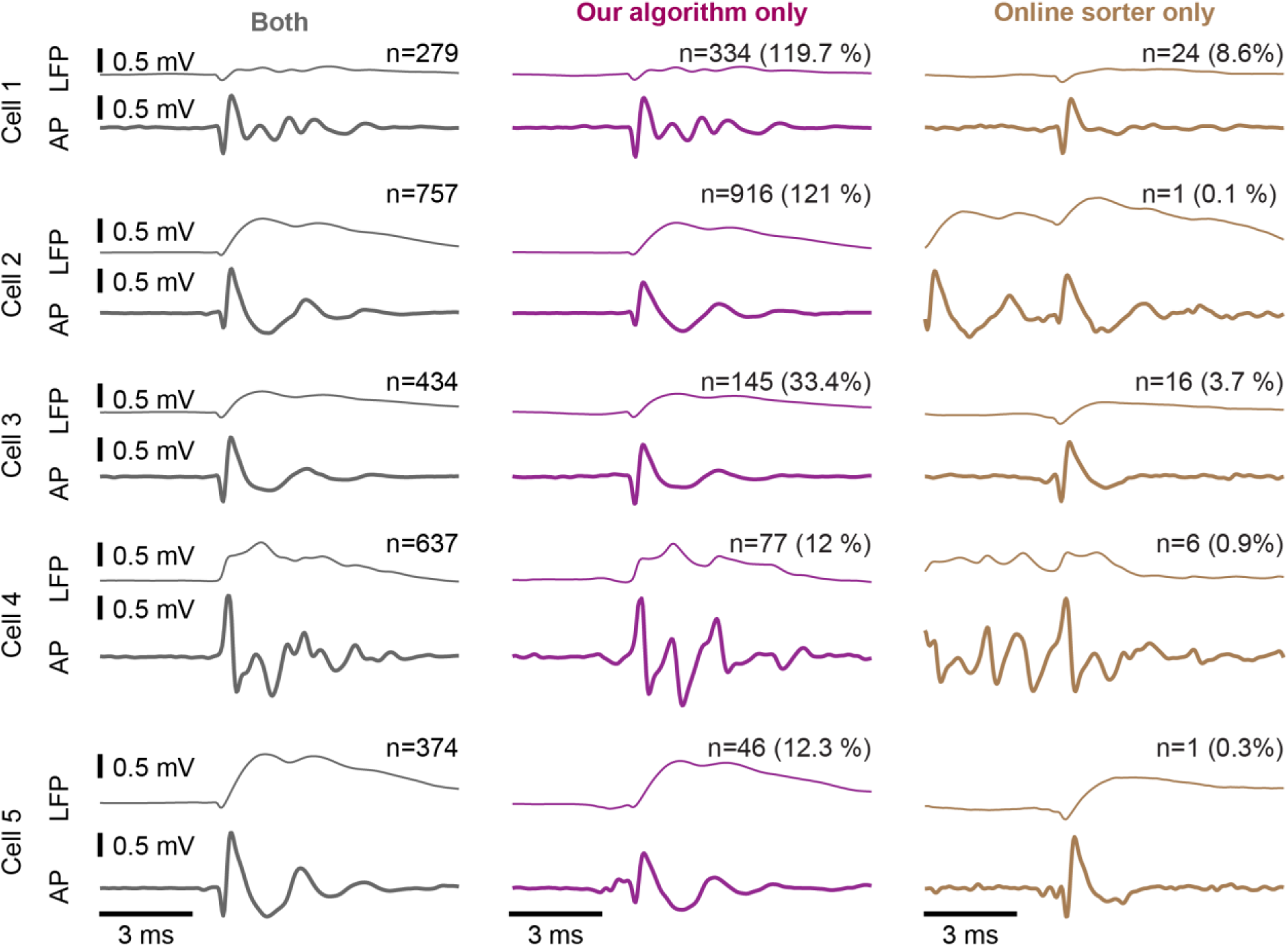
Waveforms of events labeled as CSs by our algorithm and the online sorter application MSD. Examples from seven neurons showing the average waveform in the LFP and action potentials of CSs detected by both methods (left), by our algorithm only (middle) or by the online sorter only (right).

We also evaluated to what extent the predictions from both approaches agreed with labels from a human expert. To this end, we computed the F1 score (see Methods) on short recording segments from the same 119 neurons as in the previous section for which we had “ground-truth” labels from the human expert. The F1 score is a measure of consistency in performance between an algorithm and the human expert. As shown in Fig. 6, our algorithm achieved overall higher F1 scores than the MSD, and it also showed much less variability between the different PC records (Fig. 6A). In fact, for the majority of recorded PCs, our algorithm agreed with the human expert on all CS labels, reflected by an F1 score of 1. This indicates that the predictions by our approach are more “human-like” than the ones labeled by the MSD. To achieve good performance in terms of F1 score, our algorithm also did not need a lot of training data. With only 50 training records of 232 ms of data each (sampled at 25 kHz), our algorithm outperformed the MSD algorithm (Fig. 6B). Larger training sets, of course, yielded even higher performance (Fig. 6B).

**Figure 6.**
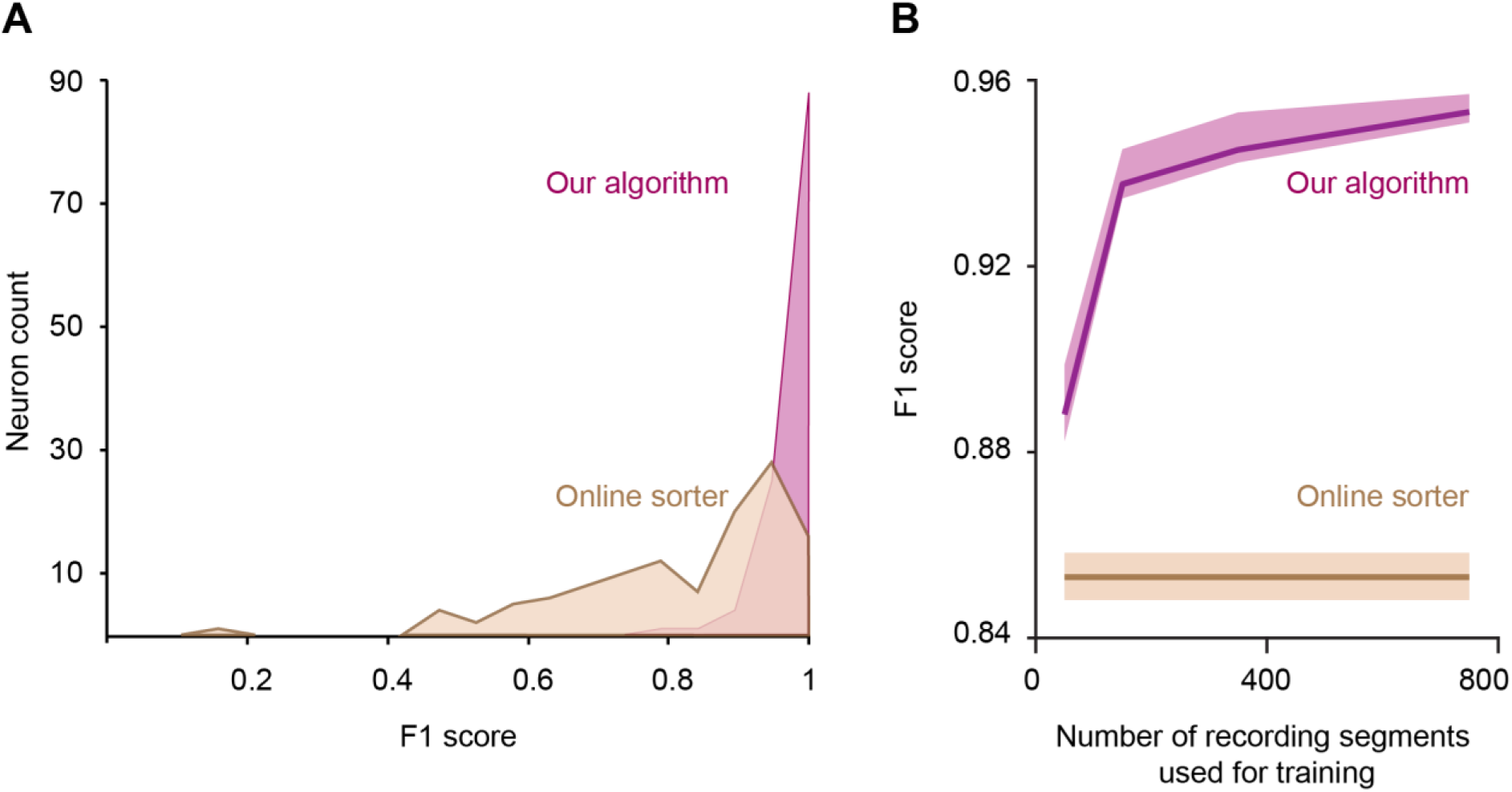
Classification agreement of our algorithm and the online sorter application MSD with a human expert. (A) Distribution of F1 scores of our algorithm and the online sorter computed by comparing CS labels with the human expert. Data from 119 neurons. (B) F1 score of our algorithm as a function of the number of recording segments used for training (pink) and F1 score achieved by the online sorter (beige). Think lines indicate the mean and the shaded area represents 95% confidence interval of the mean obtained by bootstrapping.

### CNN approach reaches human expert-level performance

Finally, for 7 PCs, we asked our human expert to fully label the entire recorded data for each neuron, instead of only a tiny training set (Methods). We then compared the CS labels of our algorithm to the ones placed by the human expert on the entire records of the neurons (spanning a time range of approximately 8-14 minutes of neural recording). Overall, the predictions of our algorithm agreed very well with the human labeling (Fig. 7A). A few events were identified as CSs by our algorithm but not by the human expert. However, also the waveforms of these events matched the waveforms of CSs that were labeled by the human expert (Fig 7A, cells 3, 5, and 6), indicating that the CSs ignored by the expert were indeed genuine CSs. For one of the PCs, the waveforms of additionally detected events indicated that our algorithm mistakenly labeled some SSs as CSs (Fig. 7, cell 7). These false positive detections, whose average predictive probability remained above the threshold (0.5) for more than 3 ms and were not removed during automatic post-processing, however, would appear as isolated clusters after dimensionality reduction (Fig. 2C). Hence, such false detections could be easily removed post-hoc by inspecting the properties of the CSs in the respective isolated cluster. For false positive labels, the average duration of pause in SS firing after these events would also be reduced to the average refractory period of SSs in this recording.

**Figure 7.**
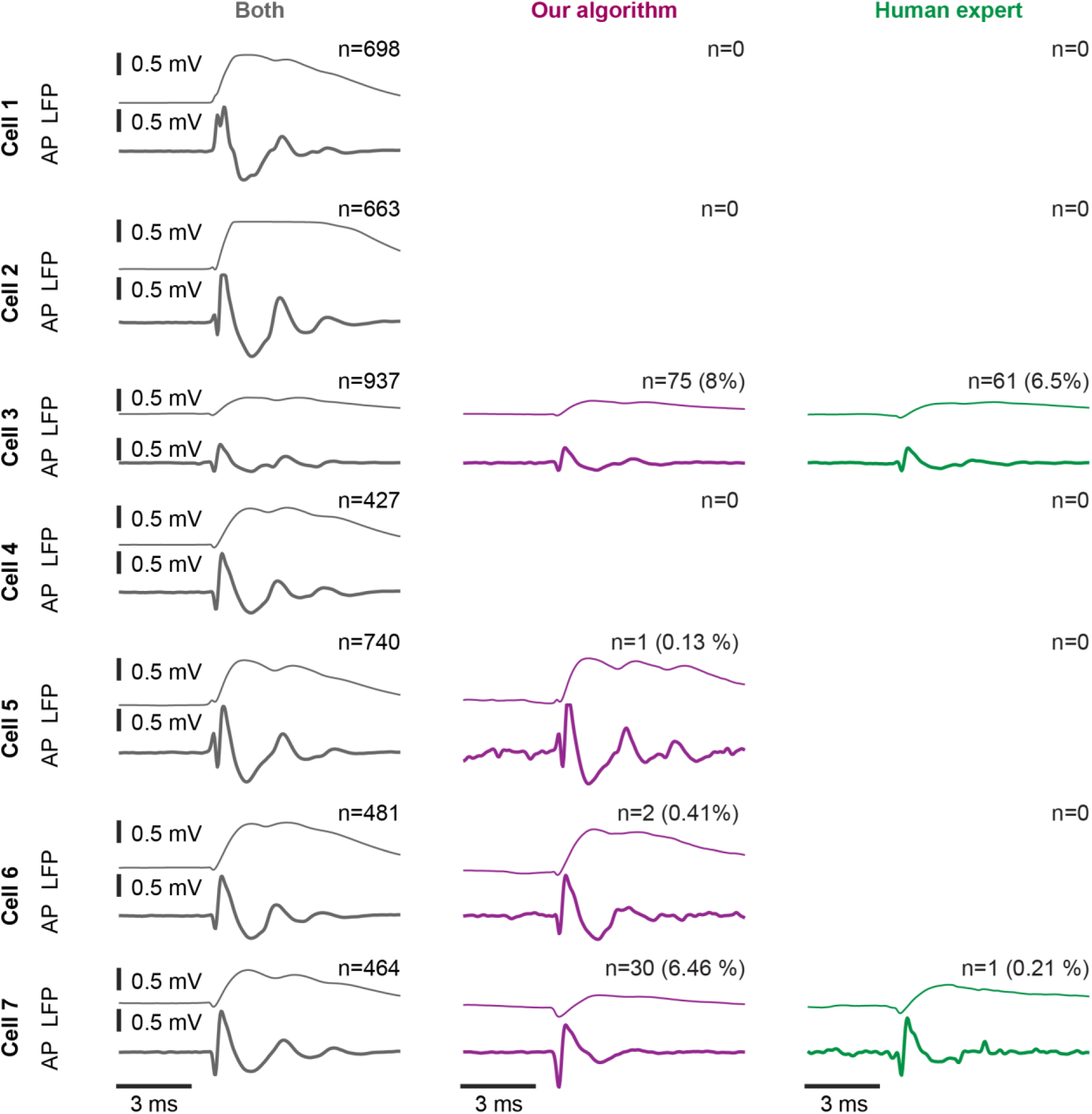
Waveforms of events labeled as CSs by our algorithm and the human expert. Examples from seven neurons showing the average waveform in the LFP and action potentials of CSs detected by both the human expert and our algorithm (left), by our algorithm only (middle) or by the human expert only (right).

The comparison with human labels further showed that our algorithm reliably identified the ends of CSs and, considering knowledge of CS start, provided a quantitative estimate of CS duration. For the recording segments from the 119 PCs, we compared the end times of all CSs that were detected by both our algorithm and the human expert. As shown in Fig. 8A, the estimate of CS end times provided by our algorithm and the human expert differed only very slightly. Correspondingly, average CS durations per neuron predicted by our algorithm and the human expert were highly correlated (ρ = 0.78, p = 1.12 × 10^−22^, Spearman correlation; Fig. 8B). In light of a possible CS duration code supplementing a CS rate code (Yang and Lisberger, 2014; Herzfeld et al., 2015; Warnaar et al., 2015; Herzfeld et al., 2018; Junker et al., 2018), it is important to precisely identify the end times of CSs and to track changes in CS duration in conjunction with behavioral changes even within individual PCs. Our algorithm was indeed capable of identifying small variations in CS duration similar to the expert. This is indicated by a strong correlation (ρ = 0.62, p = 6.81 × 10^−92^, Spearman correlation) of the residuals of human-labeled and algorithm-labeled CS end times of the selected 119 PCs, obtained by subtracting the mean CS duration of the respective PC (Fig. 8C).

**Figure 8.**
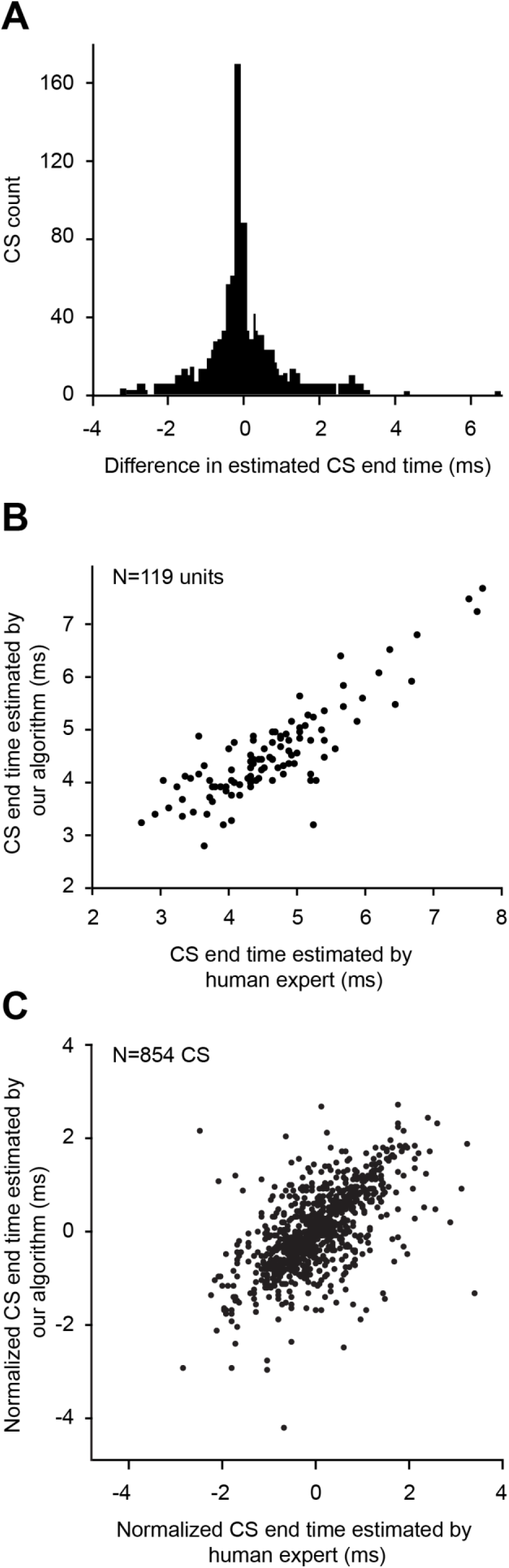
Comparison of CS end times estimated by our algorithm and by the human expert. (A) Distribution of difference in CS end times labeled by our algorithm and by the human expert. Data shows all CSs detected by both our algorithm and the human expert in short recording segments from 119 neurons. (B) Correlation of CS end times estimated by our algorithm (network) and the human expert. Each dot shows the average end time of all CSs from one neuron. (C) Correlation of all CS end times pooled across the 119 neurons. The end time of each CS was normalized by subtracting the average end time of the respective neuron.

## Discussion

This study proposes a largely automated approach to CS detection as a sensitive and reliable alternative to tedious and experience-dependent manual labeling. The approach is based on a CNN, trained on two input vectors (Fig. 9A), a high frequency band pass signal for the extraction of action potentials and a simultaneously sampled lower-frequency band pass signal reflecting LFPs. After training with surprisingly little data, our algorithm outperformed a widely used spike sorter deploying a user defined template. Moreover, our algorithm also easily caught up with the performance of an experienced human expert. Searching manually for rare events like CSs, amidst a sea of high-frequency SS signals, not only requires several weeks of tedious effort, but, as demonstrated by research on visual search (Wolfe et al., 2005; Evans et al., 2011), is also error prone, even among experts. Our network renders CS detection not just feasible, but also, more objective and systematic. Steps describing the general workflow of our algorithm are summarized in Fig. 9.

**Figure 9.**
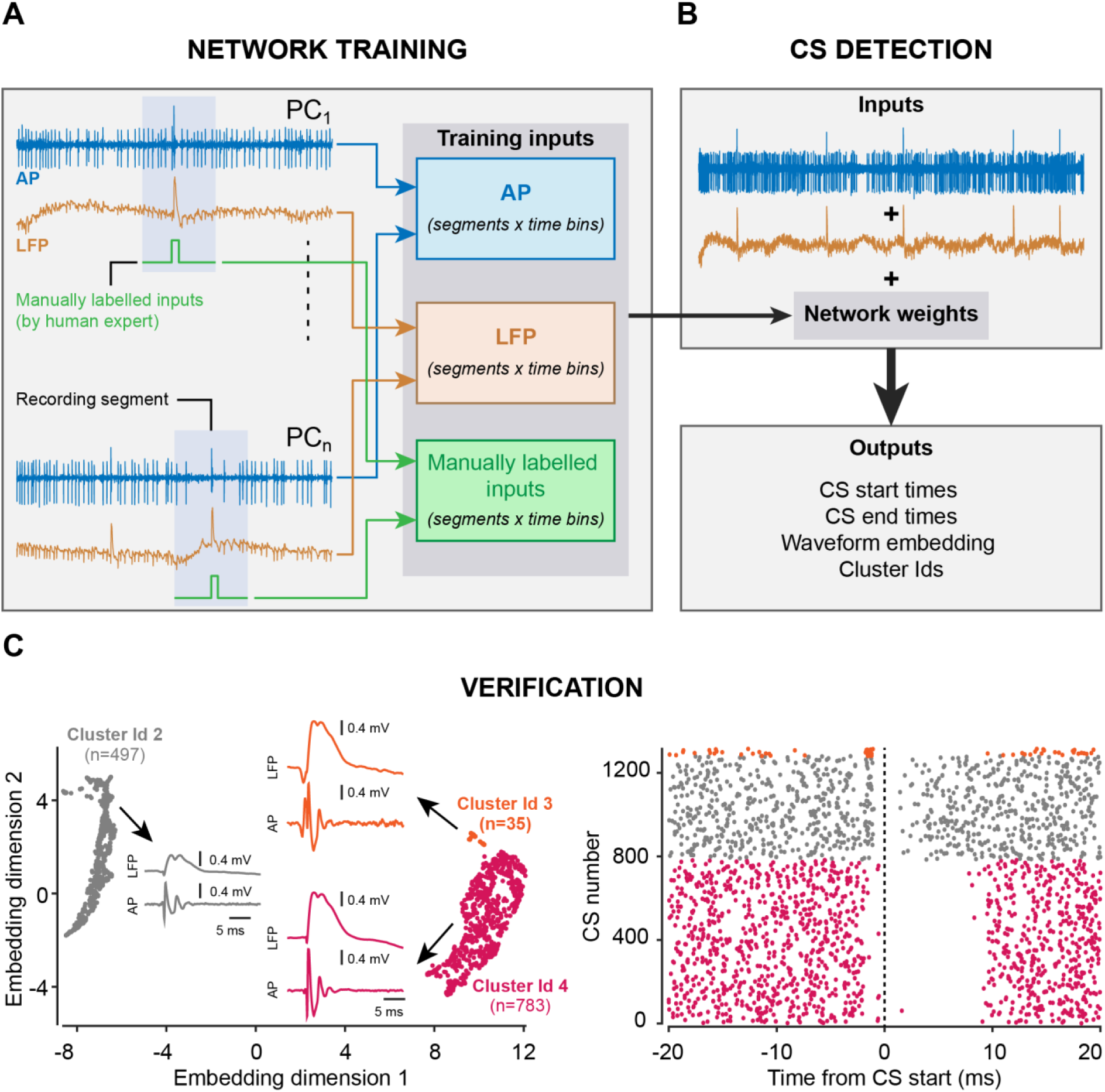
Workflow for using our algorithm. (A) The experimenter selects small segments of signal containing at least one CS each. Each segment is fed into the neural network in the form of three matrices containing the action potentials, the LFPs, and the labels separately. After training, the network outputs a set of weights. (B) The weights are used for evaluating new signals. (C) The output of the algorithm contains information about waveform shape that can be grouped in a dimensionality reduced space. This helps manual verifications, for example by inspecting the pause in SS firing rate after CS events in each cluster.

### Limitations of conventional spike sorting algorithms

The major challenge that any approach for detecting CSs meets is the polymorphic complexity of these neural events (Warnaar et al., 2015). The MSD spike sorter relies on user defined templates to identify distinct spike waveforms. However, no matter how well isolated a PC neuron may be, spike waveforms may change for internal reasons or because the position of the neuron relative to the electrode may drift over time. The MSD, like other automatic online or offline sorting approaches, tries to accommodate these changes by adapting the original template. The principal virtue of template adaptation notwithstanding, it may not be sufficient to keep track of a changing CS or, alternatively, may gradually render the template indistinguishable from the waveforms of unrelated neural activity (including the much more frequent SSs in the signal). Hence, the sorter may miss a true CS or falsely qualify other waveforms as CSs because of similar morphological features. To avoid erroneous detections and omissions, most analysts resort to manual detection. Experienced human experts may in principle reach a high level of agreement by using visual search to identify CS events. However, this approach is very tedious and therefore inevitably associated with fluctuations of attention, which jeopardizes the analyst’s performance (Wolfe et al., 2005). The tediousness of the manual detection approach is increased even further if attempts are made to pinpoint the times of CS start and end or to identify distinct features of the CS morphology such as its spikelet architecture (Warnaar et al., 2015). Conventional spike sorters based on template matching (Catz et al., 2005; Dash et al., 2010; Herzfeld et al., 2015, 2018; Junker et al., 2018) or even simpler voltage-threshold crossings can be useful to facilitate visual inspection. However, the need to double check detected CS events will forestall gains in investments of time and effort only minimally.

### Our algorithm is more sensitive and performs better than the online sorter

Our CNN-based algorithm, trained on action potential and LFP signals, clearly outperformed the MSD. Not only was it more sensitive in detecting more CSs, but it also rejected many false CSs, as compared to the MSD. This can best be seen in the example of Fig. 2C. In this figure, the Cluster 1 waveforms, despite sharing a similar shape of the initial spike component with the genuine CSs in Cluster 3, appeared as a clearly separated group in our dimensionally reduced space. These erroneous waveforms were therefore safely rejected. On the other hand, waveforms belonging to Cluster 2, neighboring the main Cluster 3, were still accepted due to close resemblance of their features to the genuine ones.

It is likely that there can be interactions between SS occurrence and CS waveform appearance. Specifically, a study on PCs in non-anaesthetized mice has demonstrated that the shape of the CS waveform can be altered by preceding SSs (Servais et al., 2004). Furthermore, recently conducted experiments on climbing fiber responses in PCs have revealed that the potassium currents, by means of voltage gating in a branch-specific manner, can regulate the climbing fiber driven calcium ion influx leading to changes in CS waveform amplitude (Zang et al., 2018). This may explain why the additional CSs detected by our algorithm might have potentially deceived the online sorter. The genuine nature of the additional CSs detected by our algorithm was confirmed with the help of another prominent physiological marker-a pause in spontaneous firing activity of SSs 10-20 ms right after the occurrence of a CS. The additional CSs that were detected by the online sorter and not by our algorithm did not show a clear suppression of SS firing.

A major factor, contributing to unsatisfactory performance of conventional sorters, is the fact that they typically rely only on information from the action potential record, rather than using complementary information from time synchronized LFP recordings, which is what human experts would do when searching PC recordings for CSs. In accordance with a very recent Principal Component Analysis (PCA) based approach (Zur and Joshua, 2019), demonstrating improved CS sorting by exploiting LFP frequency bands, the high performance of our algorithm in detecting CSs also critically relies on the use of LFP signals. The virtue of the PCA-based approach notwithstanding, it is clearly outperformed by our network. First, our approach gives a good estimate of CS occurrence without requiring a subsequent manual selection of the cluster in a principal component space. Second, as compared to the PCA, the UMAP dimensionality reduction technique is more resistant to changes in waveform shape, such as reductions in waveform amplitude due to relative shifts in position between electrode tips and cell bodies. Third, the performance of our algorithm is indifferent to occasional oscillations that may occur in the LFP signal that may impede the performance of the PCA-based approach, which relies on threshold crossings for event detection. Finally, as discussed further below, the CNN, but not the PCA, offers precise information on timing, enabling us to study CS durations much more systematically and objectively.

It is well established (Eccles et al., 1967) that each PC receives input from only one climbing fiber. Therefore, it is very unlikely to find a second CS with completely different properties in addition to the first CS in a PC record. Surprisingly, we found two PCs (see Fig. 9C for an example) for which the CNN delineated a completely separate, large cluster of CSs in addition to the main cluster. At first glance, this might have suggested a violation of the aforementioned architectural principle. However, the CSs found in the respective second clusters could be easily discarded post-hoc because of the insufficient suppression that they induced in SS firing as compared to the genuine CSs. Therefore, although rare, even if genuine CSs that belonged to a neighboring PC (Fig. 9C, seen as much smaller amplitude waveforms in Cluster 2) were captured by the electrode tip, these CSs could easily be identified based on their cluster IDs and scrutinized for selection.

To test whether our algorithm could really take over the burden of labeling CSs manually, we made a one to one comparison of the performance of the CNN and the human expert on records of 7 PCs for which all CSs had been labeled manually. Indeed, our algorithm’s performance matched the human-level expertise in detecting CSs in all PCs, except for one in which additional CSs were detected by our algorithm (Fig. 7, Cell 7). The location of these CSs in a distinct cluster in two dimensional feature space allowed the experimenter to easily evaluate the validity of the identification of the waveform as CS and, in this case, to conclude that it was spurious.

### Our algorithm detects start and end points of CSs with human-level performance

The prevailing idea of CSs serving as the “teaching-signal” for post-synaptic PCs (Marr, 1969; Albus, 1971; Ito, 1972), for which the occurrence of each CS event might be the only source of relevant information (Rushmer et al., 1976; Gellman et al., 1985), has been challenged by studies that demonstrated that the duration of action potential bursts fired by olivary neurons may vary and that this may be reflected by changes in the duration and the spikelet architecture of CSs (Llinás and Yarom, 1981; Ruigrok and Voogd, 1995; Maruta et al., 2007; Mathy et al., 2009; Bazzigaluppi et al., 2012; De Gruijl et al., 2012; Rasmussen et al., 2013; Zang et al., 2018). These observations have suggested that not only the occurrence of a CS, but also its duration may be relevant for motor learning. Addressing this possibility requires experimenters to invest even more time to manually label the start and end times of CS waveforms in addition to just detecting the events themselves. Not surprisingly, given the amount of time and effort involved, only a handful of attempts have been made to test this idea (Yang and Lisberger, 2014; Herzfeld et al., 2015, 2018; Junker et al., 2018) with inconsistent results. In order to achieve consensus, larger data sets collected under more diverse conditions would have to be explored, a necessity researchers have been reluctant to meet because of the hassles of the manual timing analysis. Since our CNN-based approach is able to effortlessly follow the performance of the human expert in detecting the start and end of the CS waveforms, by applying the expert’s “mental rules” learned during training, quantifying task related changes in the architecture of CSs collected at different times in an experiment will become much more feasible in the future.

### Deep learning as a research tool

More broadly, deep learning allows modeling non-linear relationships between input and output for which no analytical solutions may exist. It is exactly this property of deep learning that explains why this machine learning approach has recently emerged as a potentially powerful research tool, which can tremendously reduce the workload of scientists (Ciregan et al., 2012; Havaei et al., 2017; Oztel et al., 2017; Bellet et al., 2018). In light of recent developments, in which deep learning has been successfully utilized to not only design stimuli with controlled higher order statistics (Gatys et al., 2015), but also to model non-linear relationships in neural data (Ecker et al., 2018), it is not hard to imagine that the full potential of deep learning will significantly boost the pace of neuroscientific research in the coming years. Certainly, in the case of cerebellar neurophysiology, we believe that our use of deep learning to detect the rare, but relevant, CS events will allow much renewed investigation of the contentious functional roles of these events in motor control and beyond.

## Conclusion

So far, all analysis involving CSs has been based on extremely laborious, manual, or semi-automated methods lasting up to several weeks. This enormously slows down the pace of developments in the field. On the other hand, our deep learning approach can reverse this reality. For example, for a database like ours (160 PCs), our approach requires the human expert to invest only 2-3 hours of CS labeling for training purposes and another 3-4 hours to later verify the results. Given that it takes 3-4 hours to manually label all CSs found in recordings of just one PC, this investment in time is negligible compared to the alternative of manually labeling all recorded PCs. Moreover, our automated algorithm performs this task at par with human experts, and it renders more systematic valuable information about the timing and morphology of CS waveforms. The algorithm will be made available for use via an open source implementation https://github.com/jobellet/detect_CS with provisions for retraining the network to new users’ own measurements. We strongly believe that the gains in time and reliability that our tool offers may substantially facilitate the quest for a better understanding of the roles of the still largely mysterious CSs.

## Conflict of interest

The authors declare no competing financial interests

## Acknowledgements

Supported by DFG Research Unit 1847 “The Physiology of distributed computing underlying higher brain functions in non-human primates”.

